# Pyruvate kinase M1 suppresses development and progression of prostate adenocarcinoma

**DOI:** 10.1101/2021.05.09.443334

**Authors:** Shawn M. Davidson, Julia E. Heyman, James P. O’Brien, Amy C. Liu, Daniel R. Schmidt, William J. Israelsen, Talya L. Dayton, Raghav Sehgal, Roderick T. Bronson, Elizaveta Freinkman, Howard Mak, Scott Malstrom, Gary Bellinger, Arkaitz Carracedo, Pier P. Pandolfi, Kevin D. Courtney, Abhishek Jha, Ronald A. DePinho, James W. Horner, Craig J. Thomas, Lewis C. Cantley, Massimo Loda, Matthew G. Vander Heiden

**Author notes:** These authors contributed equally to this work (Julia E. Heyman, James P. O’Brien, Amy C. Liu).

## Abstract

Most cancers, including prostate cancers, express the M2 splice isoform of pyruvate kinase (*Pkm2*). This isoform can promote anabolic metabolism to support cell proliferation; however, *Pkm2* expression is dispensable for many cancers *in vivo*. Pyruvate kinase M1 (*Pkm1*) isoform expression is restricted to relatively few tissues and has been reported to promote growth of select tumors, but the role of PKM1 in cancer has been less studied. *Pkm1* is expressed in normal prostate tissue; thus, to test how differential pyruvate kinase isoform expression affects cancer initiation and progression we generated mice harboring a conditional allele of *Pkm1* and crossed this allele, as well as a *Pkm2* conditional allele, to a *Pten* loss-driven prostate cancer model. We found that *Pkm1* loss leads to Pkm2 expression and accelerates prostate cancer, while deletion of *Pkm2* leads to increased Pkm1 expression and suppresses cancer. Consistent with these data, a small molecule pyruvate kinase activator that mimics a PKM1-like state suppresses progression of established prostate tumors. PKM2 expression is retained in most human prostate cancers, arguing that pharmacological PKM2 activation may be beneficial for some prostate cancer patients.

## Introduction

Prostate cancer is the second leading cancer-related cause of death in men and given time virtually all men will develop abnormal prostate growth, including benign prostatic hyperplasia (BPH) or prostate cancer (1). Loss of the tumor suppressive lipid phosphatase *PTEN* is associated with abnormal prostate growth in both BPH and prostate cancer, with many prostate cancers exhibiting decreased *PTEN* expression due to mutation or epigenetic silencing (2-4). Loss of PTEN activity results in phosphatidylinositol (3,4,5)-triphosphate (PIP3) accumulation and activation of AKT signaling to drive uncontrolled proliferation and survival (5,6). How these signaling events promote prostate cancer have been extensively studied (7), as have the ways in which growth factor signaling pathways can affect cell metabolism (8,9). However, whether changes in metabolic enzyme expression affect prostate tumor initiation and progression is less well defined.

Changes in metabolism are necessary to sustain cancer cell proliferation (9,10). Many cancer cells exhibit increased levels of glucose uptake with elevated lactate production even in the presence of oxygen (also known as aerobic glycolysis or the “Warburg effect”)(10,11). Increased tumor glucose consumption relative to normal tissues is exploited clinically to stage cancers through utilization of ^18^FDG-PET imaging (12); however, a signal on ^18^FDG-PET can also be observed in non-cancer settings (13). Most prostate cancers grow at a slower rate than highly ^18^FDG-avid malignancies and ^18^FDG-PET is not often used in the clinical management of prostate cancer patients, leading to the notion that these cancers rely less on glucose metabolism (14). Nevertheless, prostate cancer cells metabolize glucose in culture (15,16), and ^18^FDG-PET avidity is observed in prostate cancer, including aggressive tumors (17,18). Whether changes in glucose metabolism influence prostate cancer initiation and progression has not been extensively studied.

Induction of cellular senescence can suppress cancer development (19-21). Overcoming cellular senescence is thought to be particularly important in the pathogenesis of some prostate cancers, as Pten loss results in a p53-dependent senescence response by prostate epithelial cells in preclinical models (22), and can suppress tumor formation (19,21). Changes in metabolism to increase glucose oxidation can promote senescence in some tissues (23) and senescent lesions can be hypermetabolic (24). Cellular senescence is also associated with decreased anaplerotic flux into the tricarboxylic acid cycle (25), and nucleotide deficiency can promote this phenotype (26). Taken together, these studies suggest that increased oxidative metabolism and decreased nucleotide levels can contribute to the induction or maintenance of a quiescent or senescent state; however, the relationship between metabolism and this mechanism of tumor suppression is controversial (24,27,28), and whether changes in glucose metabolism influence senescence as a tumor suppressive mechanism in prostate cancer is not known.

Regulation of pyruvate kinase activity can influence the extent of glucose oxidation and nucleotide synthesis (29-31). Most human and murine tissues express an isoform of pyruvate kinase that is encoded by the *PKM* gene (32,33). This gene produces an RNA product that is alternatively spliced to generate mRNAs encoding two different isoforms of the enzyme: PKM1 and PKM2 (34). The mRNA encoding PKM1 and PKM2 differ only in inclusion of either exon 9 for the PKM1 message or exon 10 for the PKM2 message. PKM1 is a constitutively active enzyme that promotes oxidative glucose metabolism (29,32). PKM2 is allosterically regulated, and decreased pyruvate kinase activity associated with this isoform can promote anabolic metabolism including nucleotide synthesis (32,35). Genetic deletion of the *Pkm2*-specific exon in proliferating primary mouse embryonic fibroblasts that normally express Pkm2 results in Pkm1 expression and an irreversible proliferation arrest (31). Moreover, deleting only one *Pkm2* allele also results in Pkm1 expression that leads to proliferation arrest despite continued expression of Pkm2 at wildtype levels, indicating that expression of Pkm1 rather than loss of Pkm2 is responsible for the phenotype (31). Proliferation arrest caused by Pkm1 expression can be prevented by addition of exogenous nucleotide bases (31), suggesting that high pyruvate kinase activity associated with Pkm1 expression can limit nucleotide synthesis, but whether this results in tumor suppression is not known.

While most tissues in mice express either the *Pkm1* or *Pkm2* isoform of pyruvate kinase (33), cancer cells preferentially express *Pkm2* (32,36). This is thought to be advantageous to cancer cells because Pkm2 expression allows cells to adapt pyruvate kinase activity to different cell conditions (35,37); however, why PKM2 is selected for in most cancers is controversial (33,38). Pyruvate kinase is active as a homotetramer (30,39,40). Allosteric regulators that decrease PKM2 activity function by destabilizing the active tetramer. In contrast, PKM1 is constitutively active because residues encoded by the isoform specific exon promote stable tetramer formation. Small molecule pyruvate kinase activators have been identified that stabilize the PKM2 tetramer to promote an enzyme state similar to PKM1 (30,41-43). One such PKM2 activator, TEPP-46, is bioavailable when dosed orally in mice and can inhibit xenograft growth, phenocopying the effects of PKM1 expression (30). However, *Pkm2* expression is not required for tumor growth in several mouse cancer models (33,44-50) suggesting that loss of *Pkm2* expression might limit the ability of pyruvate kinase activators to slow the growth of many cancers.

Whether high pyruvate kinase activity due to PKM1 expression is a barrier to cancer initiation is not known. PKM1 expression is reported to provide a metabolic advantage to tumors in some contexts (38), and suppress tumor growth in others (29,30,45). Because PKM1 is constitutively active, understanding where PKM1 expression suppresses tumor growth could inform which tumor types might be sensitive to pyruvate kinase activating drugs. To study how PKM1 expression affects tumor formation, we generated mice harboring a conditional allele for the unique exon included in *Pkm1*. We crossed animals harboring this *Pkm1*-conditional allele, as well as mice harboring a *Pkm2*-conditional allele (45), to a Pten loss-driven mouse prostate cancer model (22,51). We found that Pkm isoform expression profoundly impacts prostate tumor initiation and progression. Deletion of both *Pkm1* and *Pten* in the prostate results in the formation of aggressive prostate tumors that limit animal survival. In contrast, deletion of *Pkm2* and *Pten* in the prostate results in high *Pkm1* expression and suppresses tumor formation. Importantly, small molecule PKM2 activators can also suppress prostate tumor growth, and many human prostate cancers retain PKM2 expression, arguing that forcing pyruvate kinase into a high activity state might have a role in managing prostate cancer in patients.

## Results

### Pten deletion increases glucose uptake prior to formation of invasive prostate cancer

PTEN loss is sufficient to promote glucose uptake (52) even though silencing of *Pten* alone is not sufficient to transform prostate epithelial cells (53). PTEN is frequently lost in human prostate cancer (4), so to determine whether increased glucose uptake is an early consequence of Pten loss in the prostate, mice homozygous for a conditional *Pten* allele (*Pten^fl^*) were crossed to mice with a *PbCre4* allele that drives prostate-specific Cre-recombinase expression, enabling the generation of animals with prostate-restricted Pten deletion (*Ptenpc^-/-^*) (22,51,54). These mice develop prostatic intraepithelial neoplasia (PIN) at approximately 3 months of age, which can progress to invasive cancer by 6 months of age. To assess the effect of Pten loss on glucose uptake prior to the development of invasive cancers, we measured 18FDG uptake in the prostate and muscle of *Ptenpc^-/-^* mice at 7-11 weeks of age. Glucose uptake was elevated in the prostate but not the muscle of *Ptenpc^-/-^* mice (Fig. 1A), even though this time point is prior to the onset of invasive cancer (22). These findings suggest that Pten loss is sufficient to increase glucose uptake in prostate tissue, and that increased glucose uptake can occur prior to cancer initiation.

**Figure 1.**
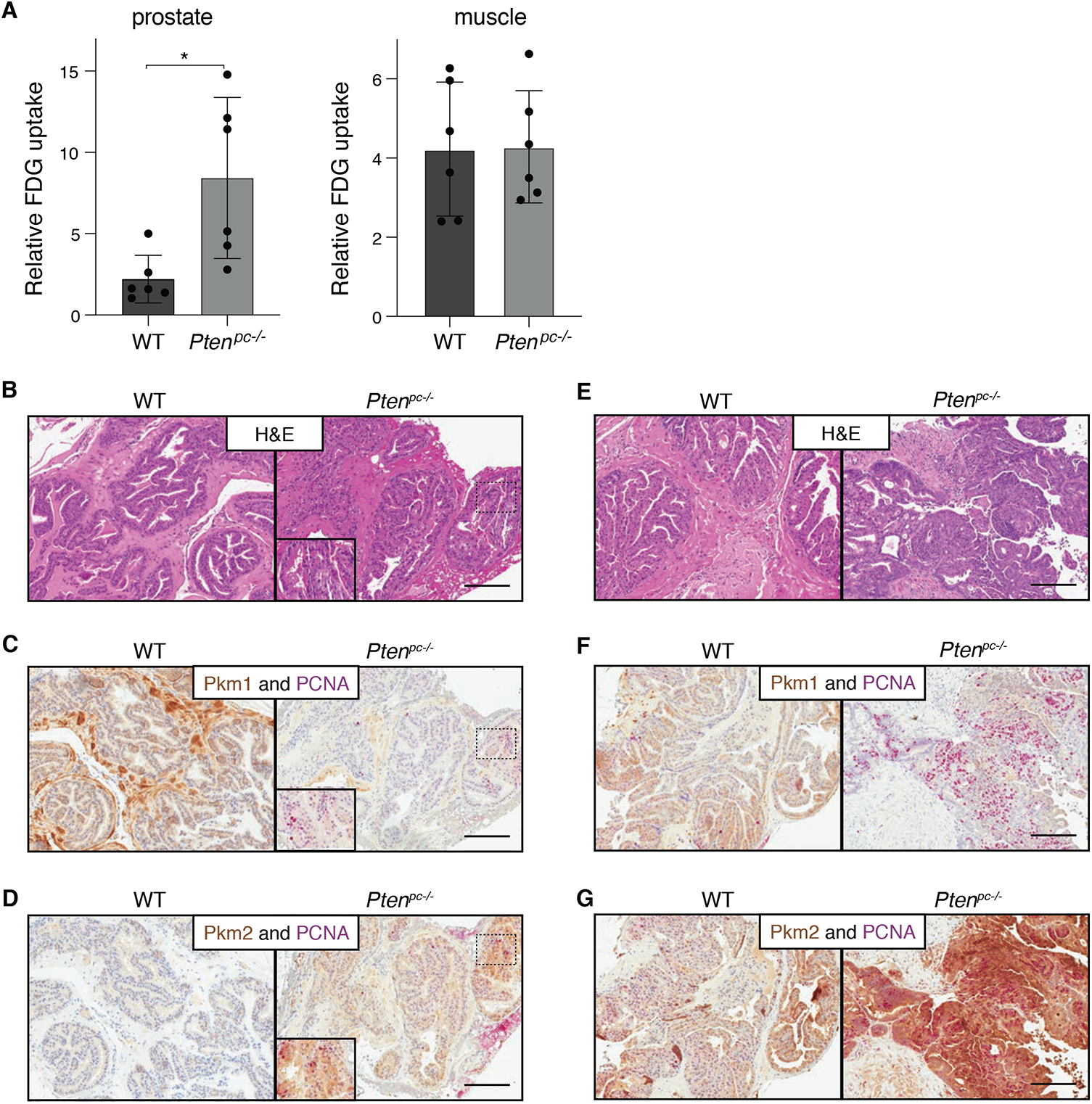
Increased glucose uptake and a change in Pkm isoform expression accompanies *Pten* loss in mouse prostate tissue. **A.** Relative [^18^F]fluoro-2-deoxyglucose (FDG) uptake into the anterior prostate and gastrocnemius muscle of 7-11 week old wild type (WT) mice and mice with prostate-specific *Pten* deletion (*Pten^pc-/-^*). Mean+/-SD is shown (n=6). The difference in FDG uptake between genotypes is significant in the prostate (*p<0.05 by Student’s t-test), but not in the muscle. **B.** H&E staining of anterior prostate tissue harvested from 3-month-old WT and *Pten^pc-/-^* mice. Scale bar = 200μm. **C.** Representative immunohistochemical staining of *Pkm1* (brown) and PCNA (pink) in anterior prostate tissue harvested from 3-month-old WT and *Pten^pc-/-^* mice. Scale bar = 200μm. **D.** Representative immunohistochemical staining of *Pkm2* (brown) and PCNA (pink) in anterior prostate tissue harvested from 3-month-old WT and *Pten^pc-/-^* mice. Scale bar = 200μm. **E.** H&E staining of anterior prostate tissue harvested from 6-month-old WT and *Pten^pc-/-^* mice. Scale bar = 200μm. **F.** Representative immunohistochemical staining of *Pkm1* (brown) and PCNA (pink) in anterior prostate tissue harvested from 6-month-old WT and *Pten^pc-/-^* mice. Scale bar = 200μm. **G.** Representative immunohistochemical staining of *Pkm2* (brown) and PCNA (pink) in anterior prostate tissue harvested from 6-month-old WT and *Pten^pc-/-^* mice. Scale bar = 200μm.

### A shift in pyruvate kinase isoform expression accompanies prostate cancer initiation

Because *Pkm1* expression is sufficient to suppress proliferation in some cells despite high glucose uptake (31), we questioned whether changes in pyruvate kinase isoform expression might be associated with cancer formation in the prostate. First, we used immunohistochemistry (IHC) and pyruvate kinase isoform-specific antibodies to determine which Pkm isoform is expressed in normal mouse prostate tissue (Fig. 1B-G, Supplementary Fig. 1). The mouse prostate has three anatomically distinct lobes (anterior prostate (AP), dorsolateral prostate (DLP), and ventral prostate (VP)) (55). In wild-type (WT) 3-month old animals, *Pkm1* is the dominant pyruvate kinase isoform expressed in the basal and luminal epithelial cells in the AP and is also prominent in the surrounding stromal cells in this lobe, with minimal *Pkm2* expression (Fig. 1B-D, Supplementary Fig. 1A), although some increase in *Pkm2* expression is observed in this lobe in 6 month old animals (Fig. 1E-G). The DLP exhibits similar epithelial *Pkm1* expression; however, low level *Pkm2* expression is also observed in luminal epithelial cells (Supplementary Fig. 1B). In contrast, the VP expresses both isoforms in epithelial cells and preferentially expresses *Pkm1* in basal cells (Supplementary Fig. 1C).

Tumor formation in *Pten^pc-/-^* mice is most prominent in the AP, and the growth of AP tumors are the major cause of mortality in this cancer model (22,56), therefore we focused on characterization of tumors arising in this lobe. By 3 months of age, when PIN is present in *Pten^pc-/-^* mice, we observed some expression of both *Pkm1* and *Pkm2* in epithelial cells, however *Pkm2* staining was more prominent (Fig. 1B-D). To assess the relationship between *Pkm1* and *Pkm2* expression and cell proliferation, we co-stained for the proliferative marker PCNA and found minimal overlap between PCNA staining and *Pkm1* expression (Fig. 1C). By 6 months of age, when invasive cancer is present in *Pten^pc-/-^* mice, there is robust *Pkm2* expression with further loss of *Pkm1* expression and increased PCNA staining (Fig. 1E-G). These data suggest that a shift from *Pkm1* to *Pkm2* isoform expression occurs with the onset and progression of *Pten*-driven prostate cancer, and that loss of *Pkm1* expression is correlated with increased cell proliferation.

### Generation of a conditional allele to prevent Pkm1 expression

To generate a conditional allele that eliminates *Pkm1* isoform expression in mouse tissues, we introduced loxP sites that flank the Pkm1-isoform specific exon 9 into the *Pkm* genomic locus of mouse embryonic stem (ES) cells using homologous recombination (Fig. 2A). Proper targeting of ES cells was confirmed by Southern blot (Supplementary Fig. 2A). Targeted ES cells were used to generate chimeric mice, which were subsequently bred to achieve germline transmission of the conditional allele and then crossed to FLP recombinase transgenic mice to delete the Neo^r^ gene. Expected targeting of the *Pkm* genomic locus was confirmed in the animals by Southern blot (Fig. 2B) and by a PCR-based approach developed for genotyping (Supplementary Fig. 2B). Intercrossing *Pkm1* conditional mice yielded progeny born in the expected Mendelian ratios that display no overt phenotypes.

**Figure 2.**
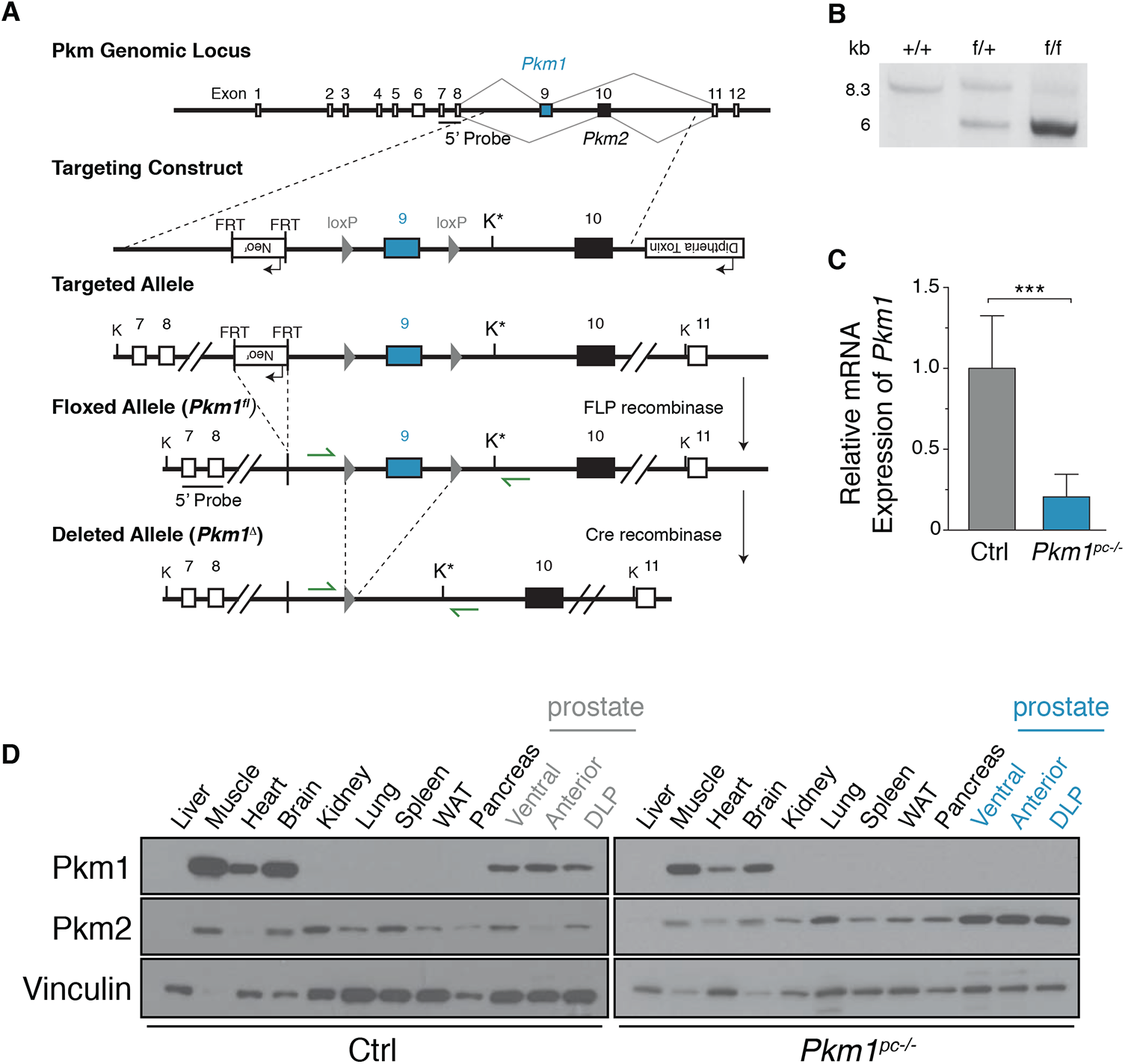
Generation and validation of *Pkm1* conditional mice. **A.** A schematic showing the mouse *Pkm* locus, construct targeting *Pkm1*-specific exon 9, and the resulting targeted, floxed, and deleted *Pkm1* alleles. The KpnI restriction enzyme sites used for Southern blot analysis are marked with “K,” and the new KpnI site introduced by the targeting vector is marked with “K*”. The location of the 5’ probe used for southern blot analysis is also indicated, as are the locations of the genotyping primers (green arrows). **B.** Southern blot analysis of KpnI-digested genomic DNA from *Pkm1*^+/+^ (+/+), *Pkm1*^+/fl^ (f/+), and *Pkm1^fl/fl^* (f/f) mice using the 5’ probe shown in (A). Digestion of genomic DNA harboring the wild-type allele (+) yields an 8.3 kb fragment, while DNA harboring the floxed allele (f) yields a ∼5.0kb fragment. **C.** *Pkm1* mRNA levels in *Pkm1^+/+^* and *Pkm1^-/-^* anterior prostate tissue as determined by quantitative RT-PCR. Mean+/-SD is shown (n=5). The difference in expression between genotypes is significant (***p<0.001 by Student’s t-test). **^D.^** Western blot analysis of *Pkm1*, *Pkm2*, and vinculin expression in various tissues including the ventral, anterior, and dorsolateral (DLP) prostate lobes that were harvested from wild type and *Pkm2^pc-/-^* mice as indicated.

To determine the effect of *Pkm1* deletion in the prostate, we crossed *Pkm1* conditional mice to animals with a *PbCre4* allele to achieve animals homozygous for the *Pkm1^fl^* allele (*Pkm1^fl//fl^ PbCre4*, hereafter *Pkm1^pc-/-^*). Examination of *Pkm1* expression in the AP from these animals showed the expected decrease in *Pkm1* mRNA transcript levels (Fig. 2C), and the absence of Pkm1 protein expression by Western blot in all prostate lobes, while Pkm1 protein expression is retained in other Pkm1-expressing tissues (Fig. 2D). Loss of Pkm1 in the prostate also resulted in increased Pkm2 expression in all three prostate lobes (Fig. 2D). These results confirm the conditional allele functions as designed, and demonstrates that deletion of *Pkm1* results in *Pkm2* expression in mouse prostate tissue in vivo.

### Pkm1 deletion promotes prostate cancer progression

To determine the effect of *Pkm1* deletion on prostate tumor initiation and progression, we crossed *Pkm1*^fl/fl^ mice to *Pten^pc-/-^* mice (hereafter, *Pkm1;Pten^pc-/-^*). We noted that *Pkm1;Pten^pc-/-^* animals developed invasive cancer more frequently in younger mice than *Pten^pc-/-^* littermates (Supplementary Fig. 3). The survival of *Pkm1;Pten^pc-/-^* animals was decreased compared with *Pten^pc-/-^* and wild-type (*Pten^pc+/+^*; WT) animals (Fig. 3A). Tumors were never observed in *Pkm1^pc-/-^* mice without *Pten* deletion when aged to 15 months, indicating that loss of *Pkm1* accelerates cancer initiated by *Pten* deletion. Furthermore, serial magnetic resonance imaging (MRI) confirmed that tumors in *Pkm1;Pten^pc-/-^* animals arose earlier and grew faster than tumors in *Pten^pc-/-^* animals (Fig. 3C, D).

**Figure 3.**
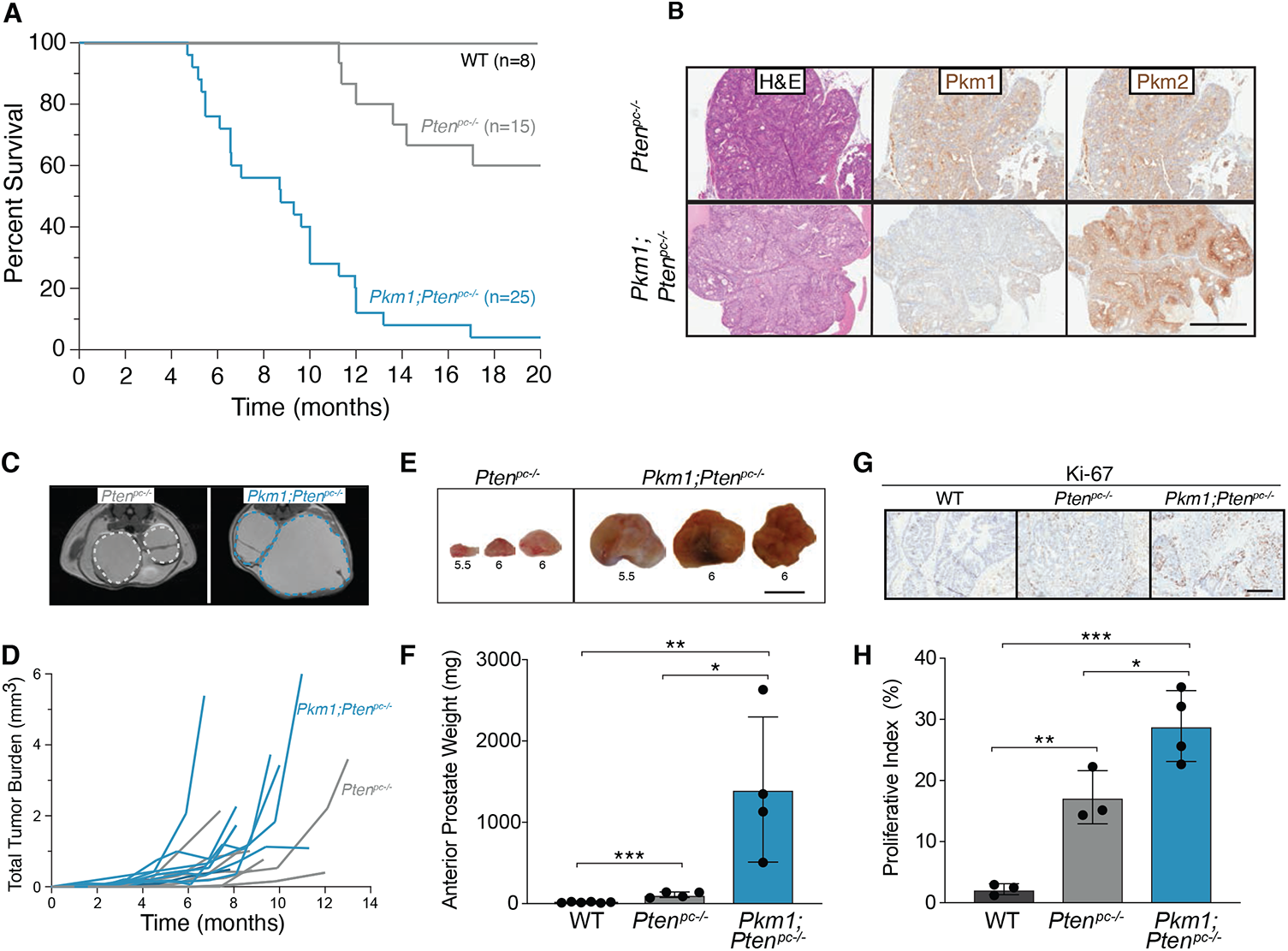
*Pkm1 deletion* together with *Pten* deletion accelerates prostate tumor growth. **A.** Kaplan-Meier curve assessing survival of a cohort of wild type (WT), *Pten^pc-/-^*, and *Pkm1;Pten^pc-/-^* mice as indicated The difference in survival between genotypes is significant (n indicated on figure, p<0.001 by log-rank test). **B.** Representative H&E and immunohistochemical staining of *Pkm1* and *Pkm2* expression in *Pten^pc-/-^*, and *Pkm1;Pten^pc-/-^* at 3 months of age. Scale bar = 500μm. **C.** Representative MRI image of 6 month old *Pten^pc-/^*^-^ and *Pkm1;Pten^pc-/-^* mice. The left and right anterior prostates are outlined in each image. **D.** Prostate tumor volume estimated from serial MRI scans over time for a cohort of *Pten^pc-/^*^-^ and *Pkm1;Pten^pc-/-^* mice as shown. Each line represents data from a single mouse, with the age of each mouse corresponding to time on the x-axis. **E.** Representative macroscopic images of anterior prostate tissue dissected from *Pten^pc-/-^* or *Pkm1;Pten^pc-/-^* mice as indicated. The age of the mouse in months at the time tissue was harvested is shown below each specimen. Scale bar 1cm. **F.** Weight of anterior prostate tissue dissected from 6 month old WT, *Pten^pc-/-^* and *Pkm1;Pten^pc-/-^* mice. Mean+/-SD is shown (WT, n=6; *Pten^pc-/-^*, n=4; and *Pkm1*;*Pten^pc-/-^*, n=4). Differences in tissue weight are significant (*p<0.05, **p<0.01, or ***p<0.001 by Student’s t-test). **G.** Representative immunohistochemical Ki-67 staining of anterior prostate tissue harvested from 6 month old WT, *Pten^pc-/-^* and *Pkm1*;*Pten^pc-/-^* mice. Scale bar = 200μm. **H.** Proliferative index assessed from Ki-67 staining of anterior prostate tissue harvested from 6 month old WT, *Pten^pc-/-^* and *Pkm1*;*Pten^pc-/-^* mice. Mean+/-SD is shown (WT, n=4; *Pten^pc-/-^*, n=3; and *Pkm1*;*Pten^pc-/-^*, n=4). Differences in proliferative index are significant (*p<0.05, **p<0.01, or ***p<0.001 by Student’s t-test).

Analysis of prostate tissue from 6-month old mice, a time where high-grade PIN and adenocarcinoma are observed in *Pten^pc-/-^* mice (22) (Supplementary Fig. 3), demonstrated that tumors from *Pkm1;Pten^pc-/-^* animals were larger than tumors from *Pten^pc-/-^* mice, and prostate size from *Pkm1;Pten^pc-/-^* was larger than that found in both *Pten^pc-/-^* and WT control mice (Fig. 3E, F). IHC for Ki-67 showed increased proliferation in both the stromal and epithelial compartments of *Pkm1;Pten^pc-/-^* tumors relative to *Pten^pc-/-^* tumors and WT prostate tissue (Fig. 3G, H), and like *Pten^pc-/-^* mouse prostate tissue, *Pkm1;Pten^pc-/-^* tumors express the androgen receptor (Supplementary Fig. 4A, B). We found no evidence of macroscopic metastases in animals of either genotype at either 6 or 12 months of age. These data argue *Pkm1* deletion accelerates growth of *Pten*-loss driven prostate tumors.

### Pkm2 deletion suppresses prostate cancer formation

To determine whether *Pkm2* is required for prostate tumor initiation and/or progression, we crossed *Pkm2*-conditional mice (*Pkm2*^fl/fl^)(45) to *Pten^pc-/-^* mice (hereafter, *Pkm2;Pten^pc-/-^*). Strikingly, abnormal prostate growth as assessed by serial MRI was only detected in older *Pkm2;Pten^pc-/-^* animals (Fig. 4A) and most of these mice lived a normal lifespan, although some ultimately developed high-grade PIN and undifferentiated tumors (Supplementary Fig. 3). Analysis of prostate tissue in 6 month old *Pkm2;Pten^pc-/-^* animals showed nearly normal appearing prostates in many animals even though all *Pten^pc-/-^* littermates developed prostate lesions by this age (Fig. 4B, Supplementary Fig. 3). We confirmed loss of Pkm2 expression in prostates from 6 month old *Pkm2;Pten^pc-/-^* mice and also observed high Pkm1 expression in tissue from these animals (Fig. 4B). Deletion of *Pkm2* had no effect on androgen receptor expression (Supplementary Fig. 4C). To assess prostate growth over time in these animals, we again used longitudinal MRI to measure prostate volumes of *Pkm2;Pten^pc-/-^* and *Pten^pc-/-^* mice. Prostates in *Pkm2;Pten^pc-/-^* mice showed minimal growth, unlike prostates of *Pten^pc-/-^* mice (Fig. 4C, D). This was confirmed at time of necropsy, where we found that prostates from ∼14-month-old *Pkm2;Pten^pc-/-^* animals were smaller and weighed less than prostate from 6 month old *Pten^pc-/-^* littermates (Fig. 4E, F). IHC for Ki-67 showed decreased proliferation in 6 month old *Pkm2;Pten^pc-/-^* prostates compared with *Pten^pc-/-^* animals (Fig. 4G, H). These data suggest *Pkm2* expression may be necessary for tumorigenesis in this tissue, and when considered together with results from *Pkm1;Pten^pc-/-^* mice, are consistent with Pkm1 expression being tumor suppressive in the prostate.

**Figure 4.**
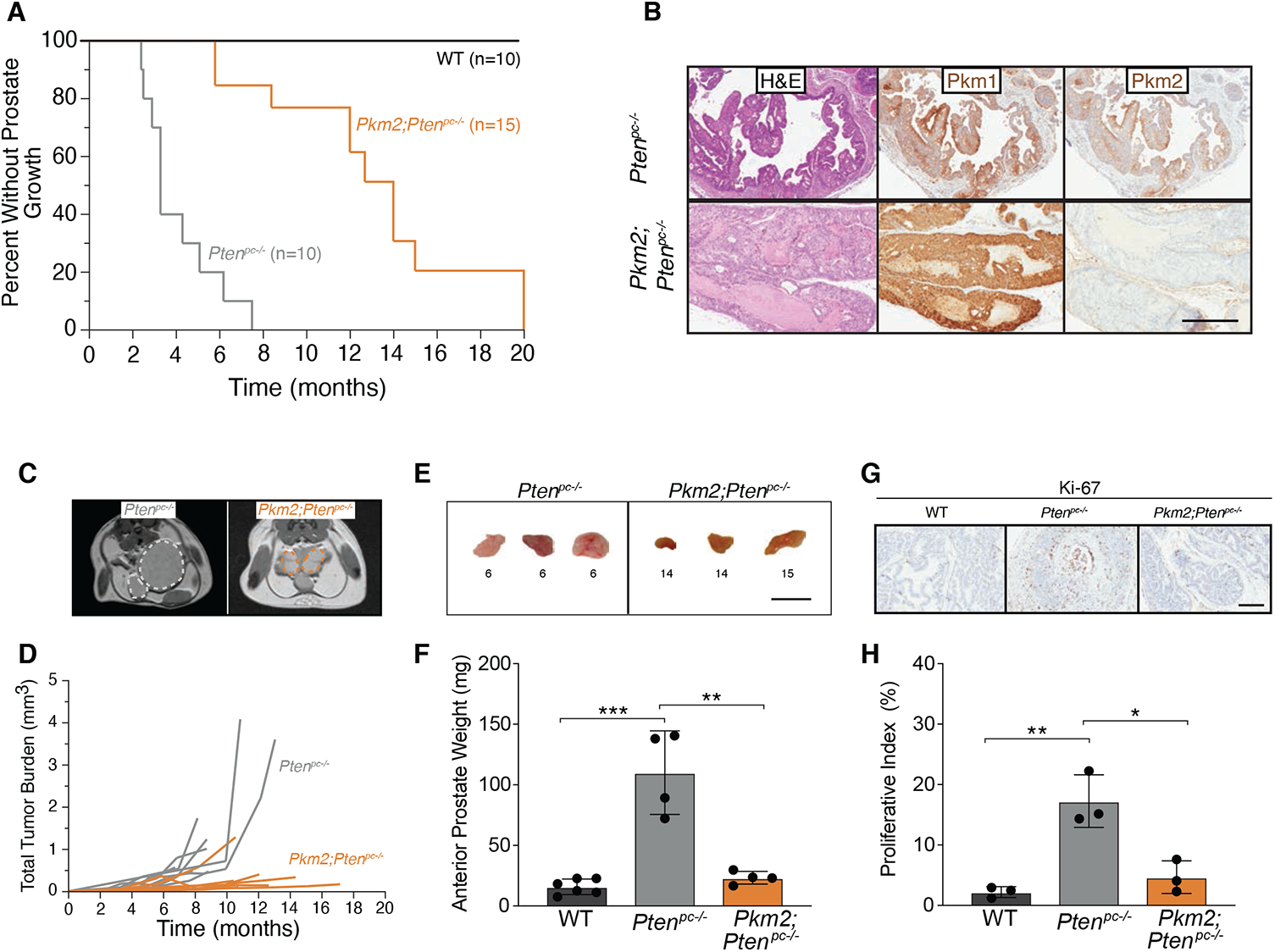
*Pkm2 deletion* suppresses prostate tumor growth induced by *Pten* deletion. **A.** Kaplan-Meier curve assessing the onset of abnormal prostate growth as assessed by MRI in wild type (WT), *Pten^pc-/-^*, and *Pkm2*;*Pten^pc-/-^* mice. The difference between genotypes is significant (n indicated on figure, p<0.001 by log-rank test). **B.** Representative H&E and immunohistochemical staining of *Pkm1* and *Pkm2* expression in *Pten^pc-/-^*, and *Pkm2;Pten^pc-/-^* at 3 months of age. Scale bar = 500μm. **C.** Representative MRI image of 6 month old *Pten^pc-/^*^-^ and *Pkm2;Pten^pc-/-^* mice. The left and right anterior prostates are outlined in each image. **D.** Prostate tumor volume estimated from serial MRI scans over time for a cohort of *Pten^pc-/^*^-^ and *Pkm2;Pten^pc-/-^* mice as shown. Each line represents data from a single mouse, with the age of each mouse corresponding to time on the x-axis. **E.** Representative macroscopic images of anterior prostate tissue dissected from *Pten^pc-/-^* or *Pkm2;Pten^pc-/-^* mice as indicated. The age of the mouse in months at the time tissue was harvested is shown below each specimen. Scale bar 1cm. **F.** Weight of anterior prostate tissue dissected from 6 month old WT and *Pten^pc-/-^* mice, and 14-15 month old *Pkm2;Pten^pc-/-^* mice. Mean+/-SD is shown (WT, n=6; *Pten^pc-/-^*, n=4; and *Pkm2;Pten^pc-/-^*, n=4). The indicated differences in tissue weight are significant (**p<0.01 or ***p<0.001 by Student’s t-test). **G.** Representative immunohistochemical Ki-67 staining of anterior prostate tissue harvested from 6 month old WT, *Pten^pc-/-^* and *Pkm2*;*Pten^pc-/-^* mice. Scale bar = 200μm. **H.** Proliferative index assessed from Ki-67 staining of anterior prostate tissue harvested from 6 month old WT, *Pten^pc-/-^* and *Pkm2*;*Pten^pc-/-^* mice. Mean +/-the SD is shown (WT, n=4; *Pten^pc-/-^*, n=3; and *Pkm2*;*Pten^pc-/-^*, n=3). The indicated differences in proliferative index are significant (*p<0.05 or **p<0.01 by Student’s t-test).

### Pkm2 deletion alters metabolism and promotes cellular senescence in Pten null prostate tissue

To assess whether pyruvate kinase isoform expression affects glucose uptake in the prostate, 6 month old WT, *Pten^pc-/-^, Pkm2;Pten^pc-/-^, and Pkm1;Pten^pc-/-^* mice were imaged with FDG-PET. Prostates from *Pkm2;Pten^pc-/-^* mice exhibited glucose uptake that was similar to prostates of WT animals, and lower than that observed in prostates from *Pten^pc-/-^* and *Pkm1;Pten^pc-/-^* mice (Fig. 5A). These data are consistent with Pkm2 loss suppressing the increased glucose uptake associated with prostate neoplasia in *Pten^pc-/-^* and *Pkm1;Pten^pc-/-^* mice.

**Figure 5.**
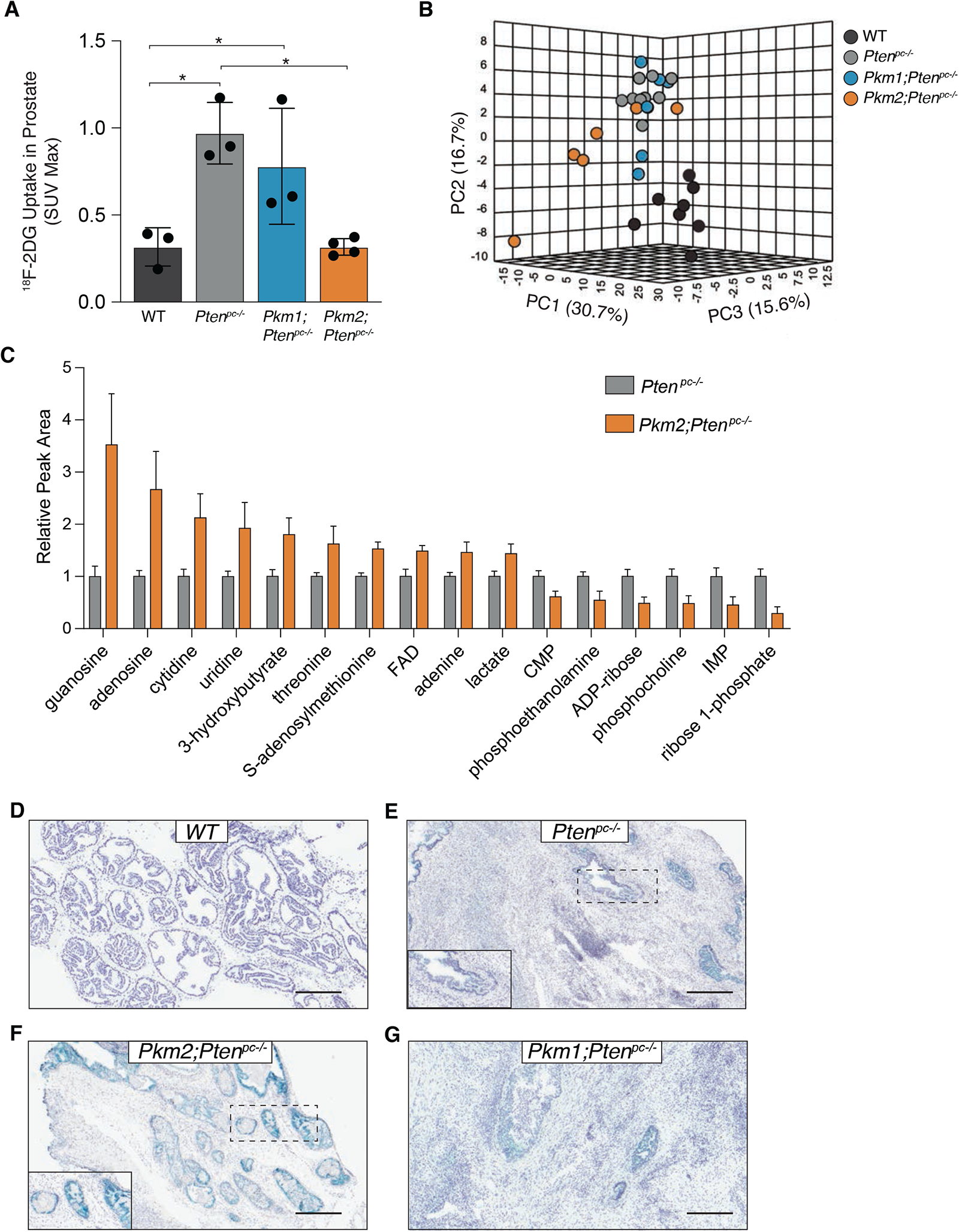
*Pkm2 deletion* suppresses increased glucose uptake, affects metabolite levels, and prolongs cellular senescence in prostate tissue following *Pten* deletion. **A.** Maximum relative [^18^F]fluoro-2-deoxyglucose signal (SUV Max) in prostate tissue as assessed by positron emission tomography of 6 month old WT, *Pten^pc-/-^, Pkm2*;*Pten^pc-/-^,* and *Pkm1*;*Pten^pc-/-^* mice. Prostate signal normalized to the intensity of the emission spectra in the heart of the same mouse is shown (WT, n=3; *Pten^pc-/-^*, n=3; *Pkm1*;*Pten^pc-/-^*, n=3; *Pkm2*;*Pten^pc-/-^*, n=4;). The indicated differences in FDG uptake are significant (*p<0.05 by Student’s t-test). **B.** Principle component analysis of 111 polar metabolites measured by LC/MS in prostate tissue harvested from 6 month old WT, *Pten^pc-/-^, Pkm1*;*Pten^pc-/-^*, and *Pkm2*;*Pten^pc-/-^* mice (WT, n=8; *Pten^pc-/-^*, n=10; *Pkm1*;*Pten^pc-/-^*, n=6; *Pkm2*;*Pten^pc-/-^*, n=6). **C.** Relative levels of all metabolites measured by LC/MS that were significantly different (p < 0.05 by Student’s t-test) in a comparison of prostate tissue harvested from 6 month old *Pten-/-* or *Pkm2;Pten^-/-^* mice (*Pten^pc-/-^*, n=10; *Pkm2*;*Pten^pc-/-^*, n=6). **D.** Representative SA-β-gal staining of anterior prostate tissue harvested from a 6 month old WT mouse. Scale bar = 500μm. **E.** Representative SA-β-gal staining of anterior prostate tissue harvested from a 6 month old *Pten^pc-/-^* mouse. Scale bar = 500μm. The area indicated by the dashed box is shown larger in the lower left inset. **F.** Representative SA-β-gal staining of anterior prostate tissue harvested from a 6 month old *Pkm2;Pten^pc-/-^* mouse Scale bar = 500μm. The area indicated by the dashed box is shown larger in the lower left inset. **G.** Representative SA-β-gal staining of anterior prostate tissue harvested from a 6 month old *Pkm1;Pten^pc-/-^* mouse Scale bar = 500μm.

In mouse embryonic fibroblasts, Pkm1-mediated suppression of cell proliferation is due to altered metabolism resulting in nucleotide depletion and impaired DNA replication (31). To determine whether changes in metabolite levels are correlated with pyruvate kinase isoform expression and/or prostate tumor growth, we examined metabolite levels in the anterior prostates from 6 month old WT, *Pten^pc-/-^, Pkm1;Pten^pc-/-^*, and *Pkm2;Pten^pc-/-^* mice (see Supplementary Data File 1 for complete dataset). Principle component analysis found that metabolite levels in WT prostate tissue are distinct from *Pten* null prostate tissue regardless of *Pkm* genotype, consistent with an effect of Pten loss on metabolism (Figure 5B). Compared to *Pten^pc-/-^ and Pkm1;Pten^pc-/-^* which clustered together, *Pkm2;Pten^pc-/-^* prostate tissue clustered separately consistent with suppression of invasive prostate cancer in these mice (Figure 5B). Interestingly, the majority of metabolites that were significantly different between *Pkm2;Pten^pc-/-^* and *Pten* null prostate tissue were related to nucleotide and redox metabolism (Supplementary Table 1, Figure 5C). Specifically, when compared to Pten null prostate tissue, *Pkm2;Pten^pc-/-^* prostate tissue exhibited increased levels of several nucleosides and decreased levels of two nucleotide monophosphates as well as ribose-1-phosphate. Elevated nucleosides may be suggestive of impaired salvage, that when coupled with decreased levels of nucleotide synthesis precursors may be indicative of impaired nucleotide metabolism in *Pkm1* expressing prostate tissue. Prostate tissue from *Pkm2;Pten^pc-/-^* mice also showed increased levels of 3-hydroxybutyrate (β-hydroxybutyrate) and lactate, potentially suggestive of a more reduced tissue redox state that would also be predicted to impair nucleotide synthesis (10,57). Of note, none of these metabolites were significantly different when comparing tissue from *Pkm1;Pten^pc-/-^* and *Pten^pc-/-^* mice (Supplementary Table 1). Taken together, these data suggest that loss of *Pkm2* with *Pkm1* expression in the setting of *Pten* loss may affect nucleotide metabolism, possibly contributing to tumor suppression.

Nucleotide depletion can underlie oncogene-induced senescence (26), and *Pten*-deletion in the prostate initially results in cellular senescence that is overcome with time, or by *Trp53* deletion (22). To evaluate whether tumor suppression by *Pkm2* loss is associated with maintenance of senescence, we examined SA-β-gal staining as a marker of senescence in prostate tissue from 6 month old WT, *Pten^pc-/-^, Pkm2;Pten^pc-/-^, and Pkm1;Pten^pc-/-^* animals. We observe increased SA-β-gal staining specifically in prostate epithelial cells in tissue from *Pten;Pkm2^pc-/-^* animals as compared to tissue from WT, *Pten^pc-/-^*, and *Pkm1;Pten^pc-/-^* mice (Fig. 5D-G). These data are consistent with a switch from *Pkm2* to *Pkm1* expression impacting tumorigenesis by promoting or maintaining *Pten* loss-induced senescence.

### Pharmacological activation of Pkm2 in Pten^pc-/-^ mice delays prostate tumor growth

Since *Pkm1* encodes a constitutively active enzyme, and low pyruvate kinase activity can promote tumor growth (40,45), we hypothesized that forcing Pkm2 into a high-activity state might suppress tumor growth in *Pten^pc-/-^* mice. To test this, we randomized *Pten^pc-/-^* mice with established tumors to treatment with TEPP-46, an orally active PKM2 activator (30), that was administered twice a day for one month. Tumor size was assessed at baseline and biweekly over the course of therapy using MRI. Most tumors from vehicle-treated *Pten^pc-/-^* mice increased in size (as defined by >50% change in tumor volume) over the one-month period of treatment, while fewer tumors from TEPP-46 treated animals grew over the same time period, with radiographic evidence of tumor shrinkage in some mice (Fig. 6, Supplementary Fig. 5). These data suggest that pharmacological activation of *Pkm2* can also suppress prostate tumor growth and supports the notion that high pyruvate kinase activity is tumor suppressive in this tissue.

**Figure 6.**
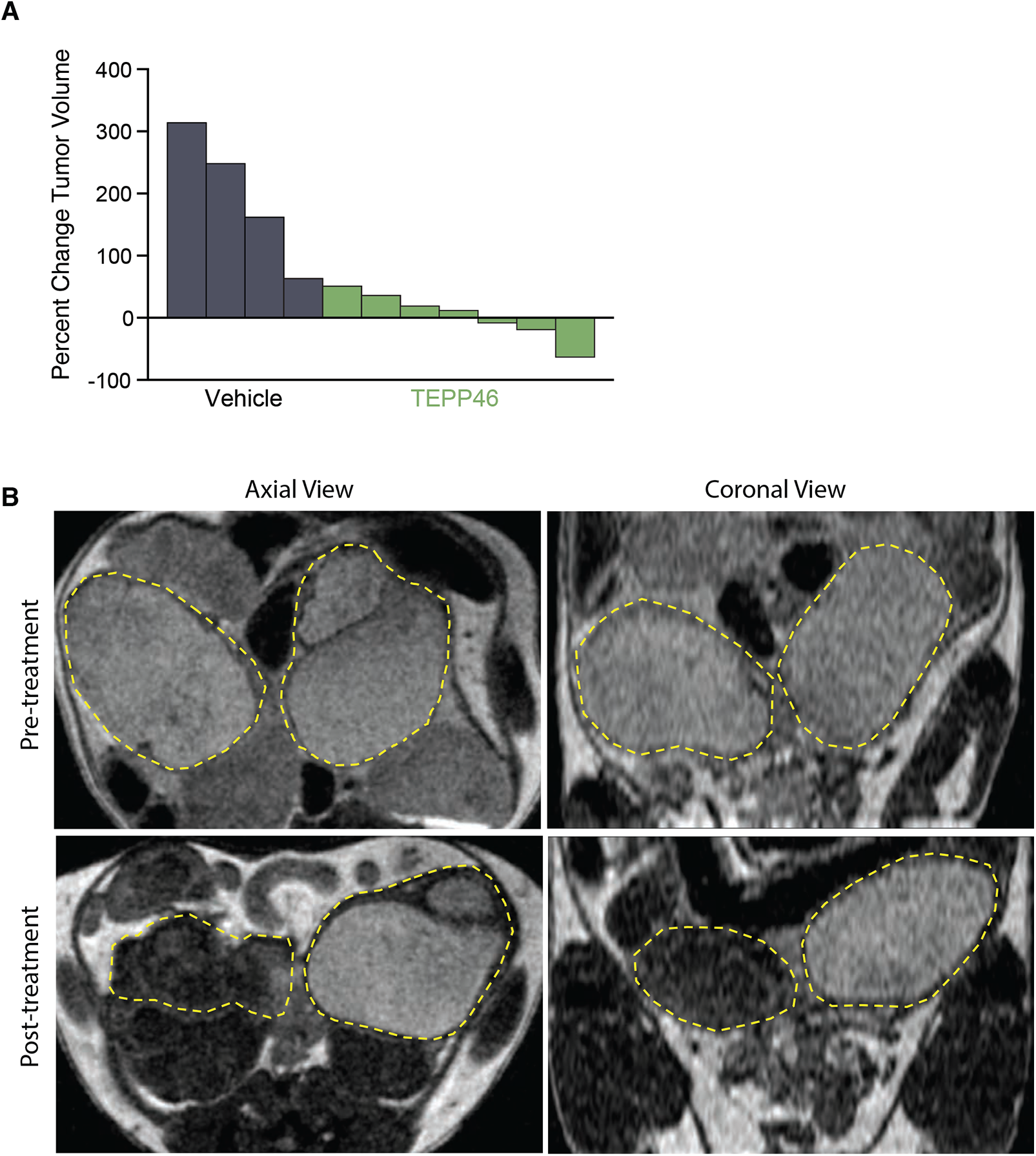
Pkm2 activator treatment reduces *Pten^pc-/-^* mouse prostate tumor growth. **A.** Waterfall plot showing the maximal change in total mouse prostate tumor volume as assessed by MRI in tumor-bearing *Pten^pc-/-^* mice after one month of twice a day treatment with vehicle or 50 mg/kg of TEPP46 as indicated (vehicle, n = 4; TEPP46, n=7). **B.** Representative MRI images from a tumor-bearing *Pten^pc-/^*^-^ mouse dosed with TEPP46 twice a day for one month. Axial and coronal view images from the same approximate anatomical plane are shown pre- and post-treatment as indicated. The left and right anterior prostates are outlined in each image.

### Clinically aggressive human prostate cancers exhibit moderate to high levels of Pkm2 expression

*Pkm2* expression is variable in many human cancers (33,45,48,50) and knowing whether this is also true in human prostate cancer is important to consider PKM2 activators as potential therapeutics. Thus, PKM2 expression was examined using IHC in sections from patients who underwent prostatectomy and in prostate cancer specimens on a prostate tumor microarray (TMA) (Fig. 7). PKM2 expression was noted in some epithelial cells in normal prostate tissue, and was retained in cancer cells in both low- and high-grade tumors, while PKM1 expression was restricted to the stromal regions of both normal and malignant prostate tissue. Of note, we observed that most human prostate tumors examined exhibit some PKM2 expression, with intermediate to high PKM2 expression found in more than half of tumors including higher Gleason grade cancers and tumors from patients with more advanced disease (Figure 7B, C). These data suggest that unlike some other human cancers, PKM2 expression is retained even in clinically aggressive prostate cancer, arguing that PKM2 activation may be effective in treating patients with this disease.

**Figure 7.**
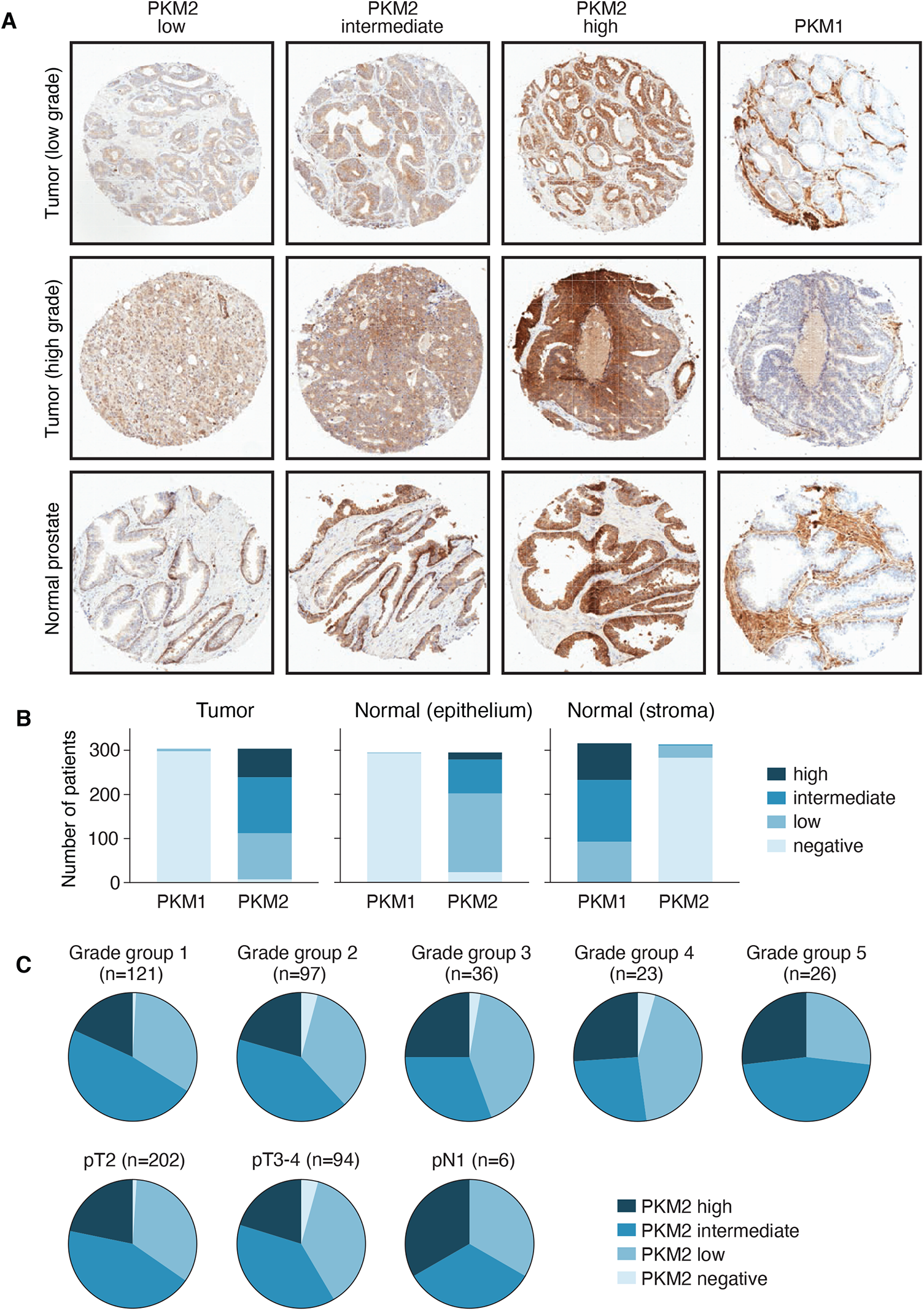
PKM1 and PKM2 expression in normal and cancerous human prostate tissue. **A.** Representative immunohistochemical staining of PKM2 and PKM1 expression in low- and high-grade human prostate tumors and normal human prostate tissue is shown. Staining scored as low intermediate, and high expression for PKM2 is indicated. The diameter of each tissue core is approximately 0.6 mm. **B.** Quantification of PKM1 and PKM2 expression by immunohistochemistry in human prostate tissue (tumor and normal) present on a tissue array containing 304 samples. Expression level is based on the scoring rubrick shown in (A). **C.** Quantification of PKM2 expression by immunohistochemistry in a human prostate cancer tissue array stratified by Gleason grade and TNM stage at the time of radical prostatectomy. The number of cases analyzed for each subset is indicated.

## Discussion

Modulation of pyruvate kinase isoform expression has large effects on *Pten* loss-driven prostate tumor initiation and growth. Despite the fact that PKM2 is the predominant isoform expressed in most human and mouse cancers, *Pkm2* expression is dispensable for the formation and growth of multiple other cancer types. Deletion of *Pkm2* in autochthonous models of breast cancer, acute myeloid leukemia, colon cancer, medulloblastoma, pancreatic cancer, hepatocellular carcinoma and sarcoma has minimal effect on cancer growth, and in some cases accelerates cancer progression (33,45-50). Thus, a tumor suppressive effect of *Pkm2* deletion in prostate tissue appears to be the exception among mouse cancer models examined to date.

In many cancer models where *Pkm2* loss does not affect tumor growth, loss of *Pkm2* is accompanied by minimal to undetectable Pkm1 expression (33,45-50). In prostate tissue, high Pkm1 expression accompanies *Pkm2* deletion. The relative correlation between Pkm1 expression and tumor suppression in various autochthonous cancer models is consistent with a tumor suppressive role for high pyruvate kinase activity. Ectopic PKM1 expression has been shown to suppress both mouse and human tumor growth in mice, even in settings where PKM2 is not deleted and there is no selective pressure to retain PKM2 (29,30,45). Nevertheless, some tumors can grow despite retaining some PKM1 expression (46,49) and a pro-tumorigenic role for PKM1 has been reported in pulmonary neuroendocrine cancers (38). However, our finding that *Pkm1* deletion in *Pten* null prostate tissue results in aggressive cancers is strongly supportive of a tumor suppressive role for Pkm1 in this organ.

The mechanism by which Pkm1 suppresses prostate tumor growth is not fully understood, although the fact that treating mice with Pkm2 activators can phenocopy Pkm1 expression suggests that high pyruvate kinase activity associated with Pkm1 is involved. One mechanism by which high pyruvate kinase activity can suppress proliferation is by affecting nucleotide synthesis (31). Nucleotide depletion can promote cellular senescence (26), and an ability to overcome senescence is a barrier to prostate cancer initiation following *Pten* loss (22). The observation that prostate cancer development involves a shift from Pkm1 to Pkm2 expression argues that a change in pyruvate kinase isoform expression may contribute to overcoming Pten loss-induced senescence.

Why high pyruvate kinase activity is particularly tumor suppressive in prostate tissue is not clear, however it could be related to the distinct metabolic phenotype of this organ. Prostate tissue synthesizes both citrate and fructose for seminal fluid (58), and either PKM1 expression or PKM2 activation can promote oxidative glucose metabolism that may promote citrate synthesis (29,30). A role for pyruvate kinase regulation in normal prostate tissue metabolism may explain in part why PKM1 is expressed in some normal prostate epithelial cells as well as explain why loss of both PKM1 and PKM2 expression is less prevalent in prostate cancer as compared to other malignancies.

The finding that Pkm2 is dispensable for tumor growth in multiple model systems, and that some cancer cells appear to proliferate despite undetectable expression of either PKM1 or PKM2 (33,44-50), suggests that the effectiveness of small molecule pyruvate kinase activators in treating cancer will be limited by loss of pyruvate kinase expression, even though molecules which activate pyruvate kinase appear to be well tolerated in both mice and humans (30,59,60). However, the finding that PKM2 is retained in the majority of human prostate cancers and that either genetic or pharmacological manipulation of pyruvate kinase appears to be tumor suppressive in an autochthonous mouse model suggests that prostate cancer might be an indication where pyruvate kinase activation could be effective for therapy. Further work is needed to test how activation of pyruvate kinase will interact with existing prostate cancer therapies, and inform how these agents should be tested in prostate cancer patients.

## Experimental Procedures

### Generation and Breeding of Pkm1 Conditional Mice and Mouse Strains

The conditional allele for *Pkm1* was generated using standard protocols to introduce loxP sites in the intronic region flanking exon 9 of the *Pkm1* gene in a manner analogous to how the *Pkm2* allele was generated (see (45). For all experiments, the *PbCre4* allele was maintained in males due to previously observed germline recombination and mosaic expression of floxed alleles when the *PbCre4* allele is transmitted through females (54). Males harboring *PbCre4, Pten*, *Pkm1*, or *Pkm2* floxed alleles were crossed to females harboring *Pten*, *Pkm1*, or *Pkm2* floxed alleles to generate prostate restricted deletion of these genes and splice products. All animals were maintained on a mixed background and littermates were used for direct comparisons.

### [^18^F]-2-Deoxyglucose Positron Emission Tomography (PET)

Animals were fasted overnight before administration of 100µCi of FDG ^18^F through a tail vein catheter. Animals were kept warm using a heated water pad and placed under 2% anesthesia during the one hour uptake time to lower background signal. Because of the small size of the prostate and its proximity to the bladder in 7-11 week old control and *Pten^pc-/-^* animals, tissue was harvested after FDG administration and gamma counts used to assess FDG uptake in prostate and gastrocnemius muscle tissue. For studies involving 6 month old mice, animals were imaged for 10 mins in PET and 1 min in CT using 720 projections at 50kv and 200µA using Sofie G8 PET/CT. Images were CT attenuation corrected and MLEM3D reconstructed. All images were decay corrected to the time of injection. The average signal intensity for three regions of each prostate was normalized to the average signal intensity for three regions in the heart of each animal.

### Southern Blot

Asp718 (Roche)-digested genomic DNA was analyzed by Southern blot using standard protocols and probe binding was visualized by autoradiography using an analogous strategy to what was described previously for the *Pkm2* allele (45). Asp718 has the same restriction site specificity as Kpn1.

### PCR Genotyping

PCR genotyping for *Pkm1* conditional mice was developed to detect and amplify the targeted *Pkm* genetic locus and performed using forward (5’-CACGCAACCATTCCAGGAGCATAT-3’) and reverse (5’-TGGTGACCTTGGCTGTCTTCCTGA-3’) primers. To genotype *PbCre4*, Forward (5’-CTGAAGAATGGGACAGGCATTG-3’) and reverse (5’-CATCACTCGTTGCATCGACC-3’) primers were used as suggested by the NCI mouse repository. Genotyping of the *Pten and Pkm2* alleles used in this study was performed as described previously (45,61).

### Western Blot and Immunohistochemistry

Western blots were performed using primary antibodies against Pkm1 (Sigma SAB4200094), Pkm2 (Cell Signaling Technology #4053), Pkm (Cell Signaling Technology #3190; Abcam ab6191), and Vinculin (Sigma, V4505, clone VIN-11-5). Fixed sections were stained with the following primary antibodies after antigen retrieval: Pkm1 (Cell Signaling Technology #7067), Pkm2 (Cell Signaling Technology #4053), PCNA (Cell Signaling Technology #2586), Ki-67 (BD Pharmingen 556003), cleaved-caspase 3 (Cell Signaling Technology #9661). Pkm1/PCNA and Pkm1/Ki-67 dual staining was quantified by scoring cells as PCNA or Ki-67 positive in a blinded fashion.

### Magnetic Resonance Imaging (MRI)

For longitudinal measurements of tumor growth, WT, *Pten^pc-/-^*, *Pten;Pkm1^pc-/-^,* or *Pten;Pkm2^pc-/-^* littermates were randomized into cohorts and prostate tissue size assessed biweekly using a Varian 7T MRI imaging system. Image sequences were acquired using the proton imaging FSEMS sequence (fast spin echo multiple slice) with TR: 4000 ms; TE: 12 ms in the axial orientation. Additional settings were as follows: 256X256 data matrix; 45X45 mm region; 1 mm thick slice; for 20 slices. OsiriX-Viewer was used for image analysis. MRI assessment of abnormal prostate growth was noted when the prostate tissue volume increased over at least two consecutive timepoints.

### β-Galactosidase Senescence Staining of Tissues

β-galactosidase staining was conducted as previously reported (62). In brief, fresh frozen-sections were cut to 8µm thickness and briefly fixed in paraformaldehyde. A solution containing one milligram of 5-bromo-4-chloro-3-indoyl β-D-galactoside (X-gal) per mL (diluted from a stock of 20mg of dimethylformamide per mL) with 40 mM citric acid/sodium phosphate pH 5.5, 5 mM potassium ferricyanide in 150 mM NaCl_2_ and 2 mM MgCl_2_ was applied to the tissue. Sections were incubated in a CO_2_ free incubator at 37°C for 12-16 hours and then visualized by conventional light microscopy.

### Metabolite Measurement and Analysis

For metabolite extraction, 10-40mg of anterior prostate tissue was weighed and homogenized cryogenically (Retsch Cryomill) prior to extraction in chloroform:methanol:water (400:600:300). Samples were centrifuged to separate aqueous and organic layers, and polar metabolites were dried under nitrogen gas for subsequent analysis by mass-spectrometry. For liquid chromatography mass spectrometry (LC-MS), dried metabolites were resuspended in water based on tissue weight, and valine-D8 was used as an injection control (63). LC-MS analyses were conducted on a QExactive benchtop orbitrap mass spectrometer equipped with an Ion Max source and a HESI II probe, which was coupled to a Dionex UltiMate 3000 UPLC system (Thermo Fisher Scientific, San Jose, CA). External mass calibration was performed using the standard calibration mixture every seven days. Sample was injected onto a ZIC-pHILIC 2.1 × 150 mm (5 µm particle size) column (EMD Millipore). Buffer A was 20 mM ammonium carbonate, 0.1% ammonium hydroxide; buffer B was acetonitrile. The chromatographic gradient was run at a flow rate of 0.150 ml/min as follows: 0–20 min.: linear gradient from 80% to 20% B; 20–20.5 min.: linear gradient from 20% to 80% B; 20.5–28 min.: hold at 80% B. The mass spectrometer was operated in full-scan, polarity switching mode with the spray voltage set to 3.0 kV, the heated capillary held at 275°C, and the HESI probe held at 350°C. The sheath gas flow was set to 40 units, the auxiliary gas flow was set to 15 units, and the sweep gas flow was set to 1 unit. The MS data acquisition was performed in a range of 70–1000 m/z, with the resolution set at 70,000, the AGC target at 106, and the maximum injection time at 80 msec. Relative quantitation of polar metabolites was performed with XCalibur QuanBrowser 2.2 (Thermo Fisher Scientific) using a 5 ppm mass tolerance and referencing an in-house library of chemical standards.

PCA analysis was performed using MetaboAnalyst 4.0 (McGill University). The input dataset included 111 metabolites that were detectable in primary prostate tumors of all 3 genotypes and wild-type prostate (4 groups total). The dataset was filtered for metabolites with near-constant values by IQR. Missing values (0.7%) were replaced with a value equal to half of the minimum value detected (assumed to be the detection limit) for a given metabolite. Peak areas were log transformed and centered at the mean.

### TEPP-46 Treatment of Pten^pc-/-^ Mice

6 month old *Pten^pc-/-^* animals were serially imaged using MRI until tumors were estimated to be >2mm^3^, and then randomized to receive either *Pkm2* activator (TEPP-46) or a vehicle control delivered twice daily via oral gavage at a final dose of 50 mg/kg in a volume less than 250µL for 4 weeks. The ages of mice treated varied from 8-18 months with cohorts selected based on tumor size. TEPP-46 was formulated in 0.5% carboxymethyl cellulose with 0.1% v/v Tween 80 as previously reported(30).

### Analysis of Human Prostate Cancer

*PKM1* and *PKM2* expression was determined by IHC in clinically annotated human prostate cancer tissue sections collected at Dana-Farber Cancer Institute during routine clinical care. Collection of tissue was approved by the Institutional Review Board of the Dana Farber Cancer Institute (Protocol 01-045) and Partners Healthcare (IRB 2006P000139). Briefly, formalin-fixed, paraffin-embedded (FFPE) prostate tissue from 345 patients (including 3 tumor cores and 2 matched normal cores per patient) were arrayed on seven panels. Tissue microarray (TMA) H&E sections were reviewed by a board-certified genitourinary pathologist (ML) to confirm presence of tumor and normal prostate. Corresponding unstained TMA sections were stained with antibodies that detect either PKM1- or PKM2-specific epitopes as described above. In total, 304 patients for which adequate tumor or normal tissue were present were included in the final analysis. Each core was scored for PKM1 and PKM2 and assigned a categorical variable (negative, low, intermediate, or high) based on the intensity of staining. Each individual patient was assigned the median category of the 3 tumor cores (expression in tumor) and higher category of the 2 normal cores (expression in normal prostate).

### Statistical Analysis

Log-rank tests were performed to determine significance in survival or tumor incidence (SPSS Statistics). Two-tailed paired and unpaired Student’s T-test were performed for all other experiments unless otherwise specified (GraphPad PRISM 7). Results for independent experiments are presented as mean ± SEM; results for technical replicates are presented as mean ± SD.

## Acknowledgements

We thank the Swanson Biotechnology Center for tissue processing and members of the Vander Heiden Laboratory for thoughtful discussions. S.M.D. was supported by an NSF Graduate Research Fellowship and T32GM007287. L.F. acknowledges support from W81XWH-15-1-0337 from the Department of Defense. D.R.S. acknowledges support by the Joint Center for Radiation Therapy Foundation and the Harvard University KL2/Catalyst Medical Research Investigator Training award (TR002542). C.J.T. acknowledges support from the Division of Preclinical Innovation, National Center for Advancing Translational Research and the Center for Cancer Research, National Cancer Institute. L.C.C. acknowledges support from R35CA197588. M.G.V.H acknowledges support from the Ludwig Center at MIT, the Burroughs Wellcome Fund, the Damon Runyon Cancer Research Foundation, the MIT Center for Precision Cancer Medicine, Stand up to Cancer, the Emerald Foundation, the NIH (P30CA1405141, R35CA242379, R01CA168653, K08CA136983, P50CA090381), and a faculty scholar award from the Howard Hughes Medical Institute.

## Declarations

All animal experiments were approved by the MIT Committee on Animal Care. All tissue analyzed from prostate cancer patients was obtained via protocols approved by the Institutional Review Board of the Dana Farber Cancer Institute (Protocol 01-045) and Partners Healthcare (IRB 2006P000139).

## Author contributions

Conceptualization: S.M.D., M.G.V.H.

Methodology: S.M.D., J.E.H., J.P.O, A.C.L., D.R.S., W.J.I., T.L.D., R.S., E.F., H.M., S.M., G.B., A.C., P.P.P., K.D.C., J.F., A.J., J.W.H., C.J.T, M.L.

Formal Analysis: S.M.D., R.T.B., D.R.S.

Investigation: S.M.D., M.G.V.H

Writing: Original Draft, S.M.D., M.G.V.H.

Writing: Review & Editing, S.M.D., J.E.H., J.P.O., A.C.L., D.R.S., M.G.V.H.

Funding Acquisition: S.M.D., R.A.D., L.C.C., M.G.V.H.

Resources: L.C.C., M.G.V.H.

Supervision, M.G.V.H.

## Supplementary Information

### Supplementary Figure and Table Legends

**Supplementary Figure 1.**
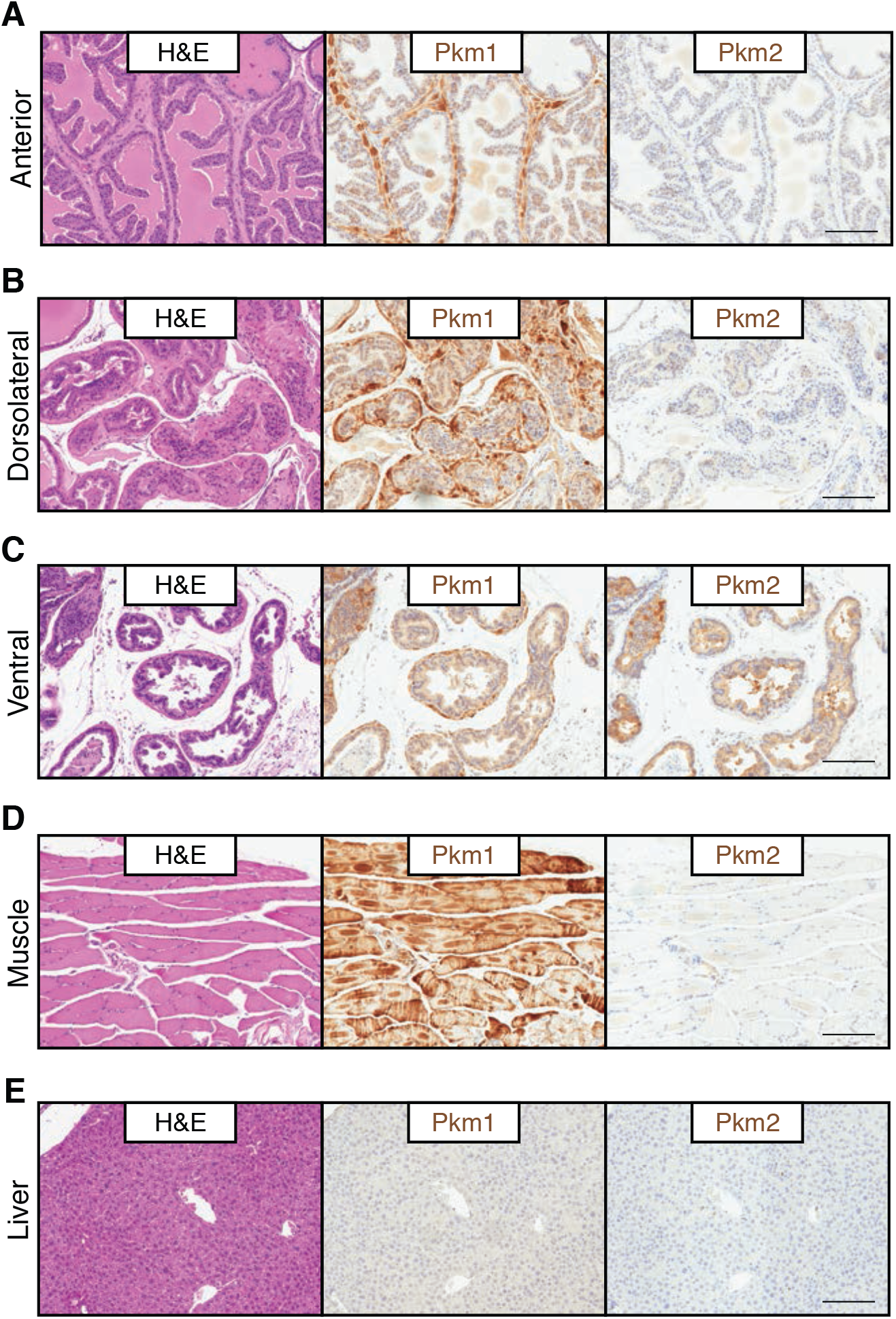
Pkm isoform expression in murine prostate tissue. **A.** Representative H&E staining of, as well as representative immunohistochemical staining of Pkm1 or Pkm2 expression in, anterior prostate tissue harvested from 3-month-old wild type mice as indicated. Scale bar = 200μm. **B.** Representative H&E staining of, as well as representative immunohistochemical staining of Pkm1 or Pkm2 expression in, dorsolateral prostate tissue harvested from 3-month-old wild type mice as indicated. Scale bar = 200μm. **C.** Representative H&E staining of, as well as representative immunohistochemical staining of Pkm1 or Pkm2 expression in, ventral prostate tissue harvested from 3-month-old wild type mice as indicated. Scale bar = 200μm. **D.** Representative H&E staining of, as well as representative immunohistochemical staining of Pkm1 or Pkm2 expression in, quadriceps muscle tissue harvested from 3-month-old wild type mice as indicated. Scale bar = 200μm. Muscle tissue is known to express PKM1 and serves as a positive control. **E.** Representative H&E staining of, as well as representative immunohistochemical staining of Pkm1 or Pkm2 expression in, liver tissue harvested from 3-month-old wild type mice as indicated. Scale bar = 200μm. Liver tissue expresses neither PKM1 nor PKM2 and serves as a negative control.

**Supplementary Figure 2.**
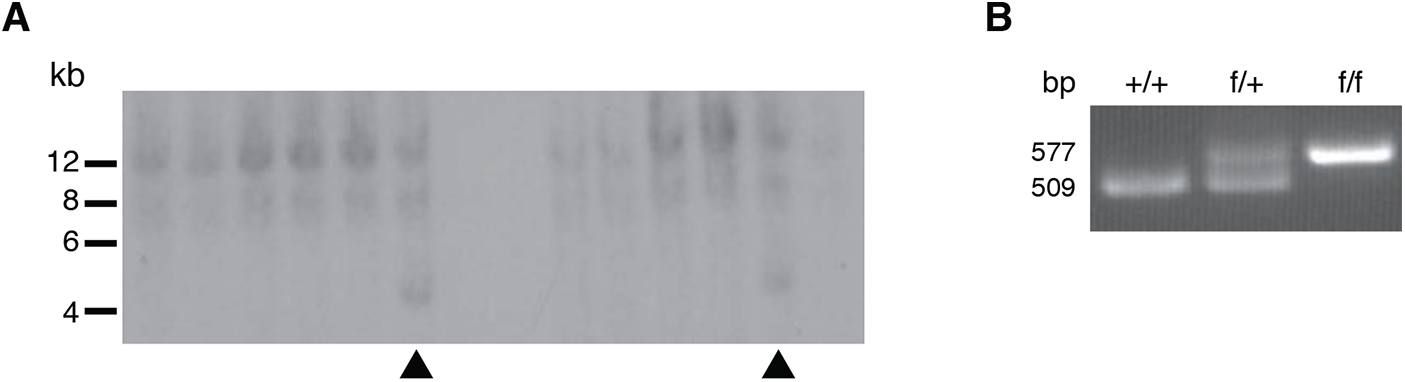
Pkm1 conditional allele characterization. **A.** Southern blot screening of embryonic stem cell clones for homologous recombination of the *Pkm1* targeting construct using the 5’ probe indicated in Figure 2A. Insertion of a novel KpnI site results in a new ∼5.0 kb fragment from the targeted locus following digestion of genomic DNA. Two clones with successful integration are marked with an arrowhead. **B.** PCR genotyping of genomic DNA from *Pkm1*^+/+^ (+/+), *Pkm1*^fl/+^ (f/+), and *Pkm1^fl/fl^* (f/f) mice as indicated. Genotyping primers anneal at sites indicated in Fig. 2A to produce amplicons of 509 bp from the Pkm wild-type allele (*Pkm^+^*) and 577 bp from the conditional Pkm allele with flanking LoxP sites (*Pkm1^fl^*).

**Supplementary Figure 3.**
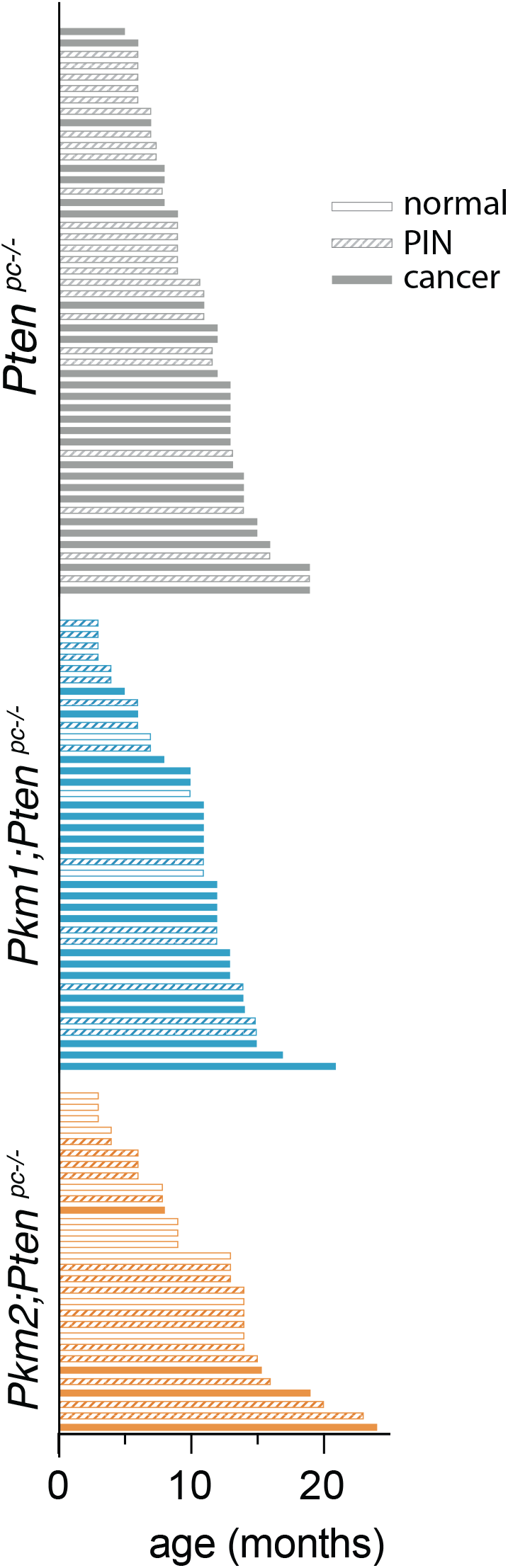
Impact of Pkm1 or Pkm2 loss on Pten-loss driven prostate neoplasia. Swimmers plot summarizing results from histopathological analysis of prostate tissue harvested from *Pten^pc-/-^, Pkm2*;*Pten^pc-/-^,* and *Pkm1*;*Pten^pc-/-^* mice at the indicated ages. Open bars represent mice with no evidence of neoplasia, hatched bars represent mice with PIN, and filled bars represent mice with invasive cancer.

**Supplementary Figure 4.**
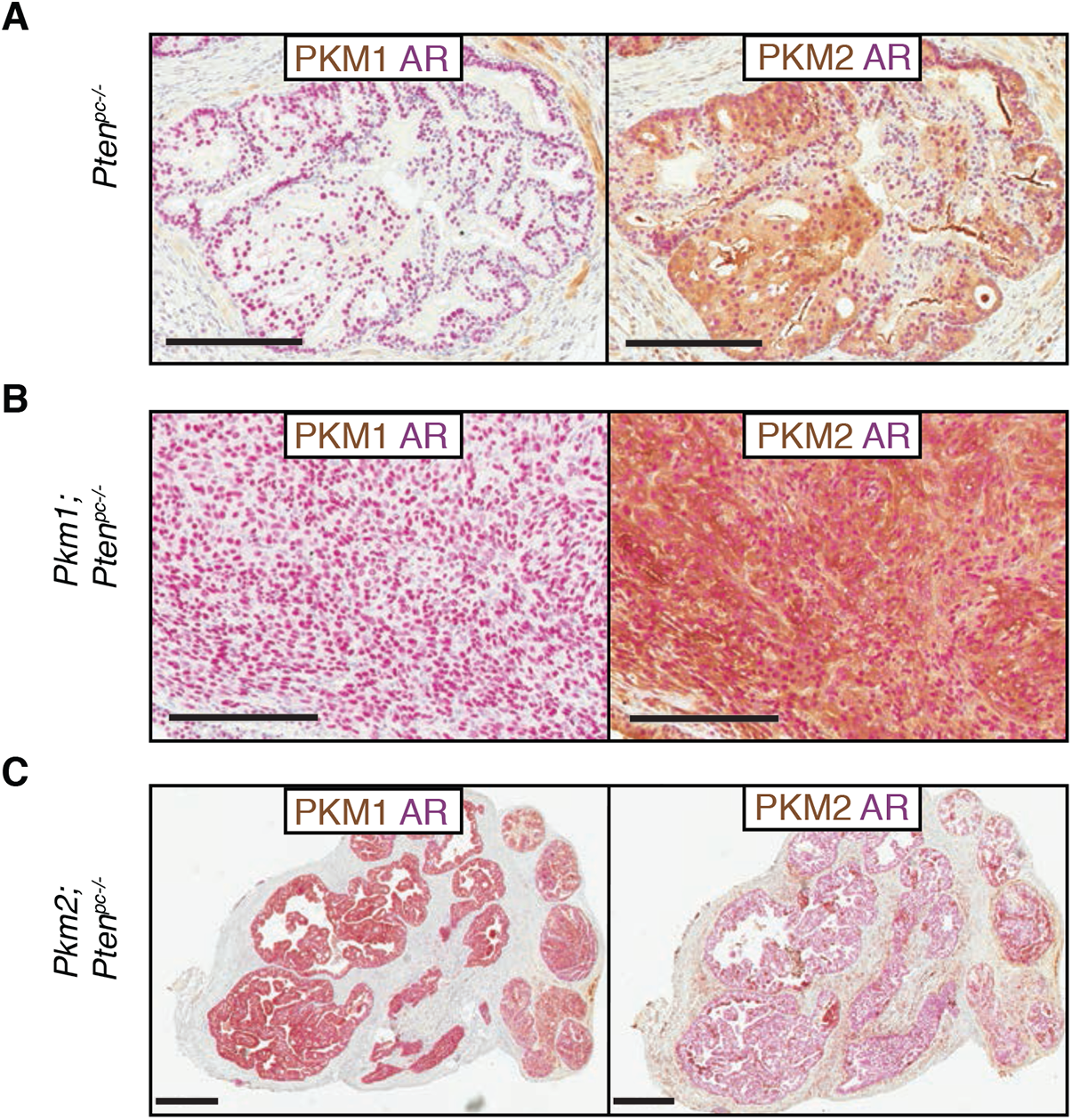
Androgen receptor expression is not affected by pyruvate kinase isoform expression in Pten-loss driven prostate lesions. **A.** Representative immunohistochemical staining of *Pkm1* (brown) and the androgen receptor (AR, pink), or *Pkm2* (brown) and the androgen receptor (AR, pink) in anterior prostate tissue harvested from 6-month-old *Pten^pc-/-^* mice as indicated. Scale bar = 200μm. **B.** Representative immunohistochemical staining of *Pkm1* (brown) and the androgen receptor (AR, pink), or *Pkm2* (brown) and the androgen receptor (AR, pink) in anterior prostate tissue harvested from 6-month-old *Pkm1*;*Pten^pc-/-^* mice as indicated. Scale bar = 200μm. **C.** Representative immunohistochemical staining of *Pkm1* (brown) and the androgen receptor (AR, pink), or *Pkm2* (brown) and the androgen receptor (AR, pink) in anterior prostate tissue harvested from 6-month-old *Pkm2*;*Pten^pc-/-^* mice as indicated. Scale bar = 500μm.

**Supplementary Figure 5.**
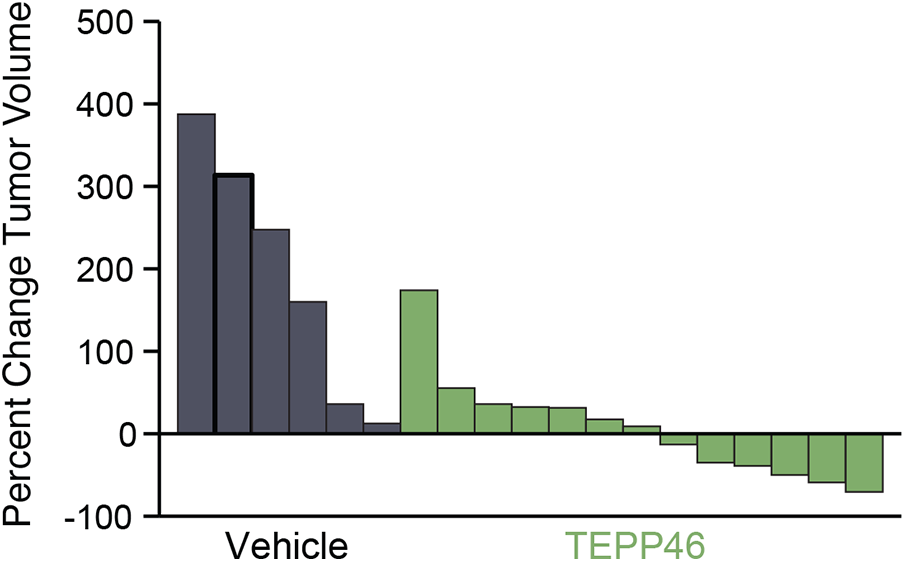
Pkm2 activator treatment reduces *Pten^pc-/-^* mouse prostate tumor growth. Waterfall plot showing the maximal change in prostate tumor volume as assessed by MRI for *Pten^pc-/-^* mice treated with vehicle or 50 mg/kg of TEPP46 as indicated. These data are from the same study presented in Fig. 6A, but display the max change in tumor volume present in each prostate lobe.

**Supplementary Table 1.**
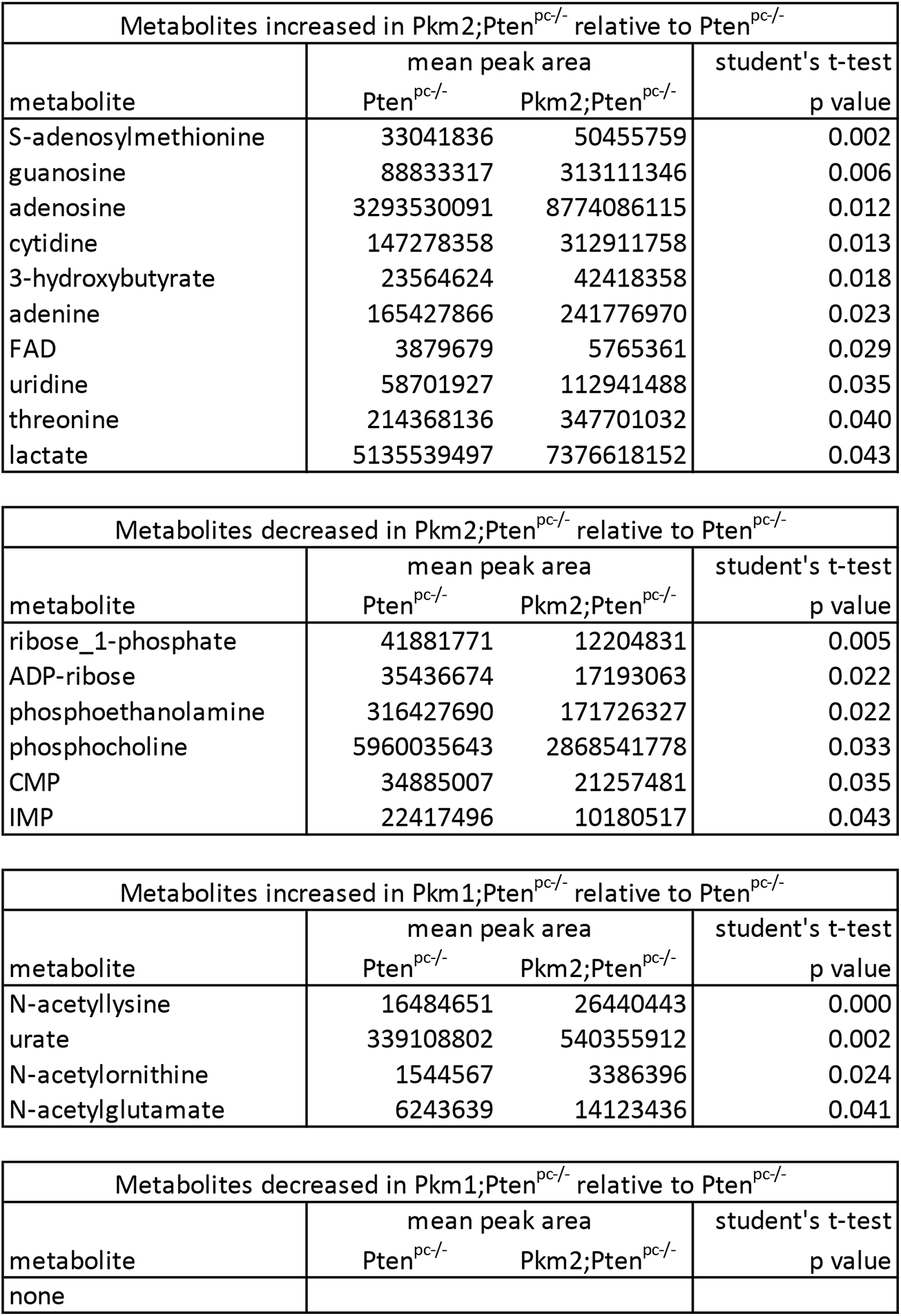
Impact of pyruvate kinase isoform expression on metabolite levels in prostate tissue from *Pten^pc-/-^* mice. Relative levels (mean peak area) for all metabolites measured by LC/MS that were significantly different (p<0.05 by Student’s t-test) in a comparison of prostate tissue harvested from 6 month old *Pkm2*;*Pten^pc-/-^* and *Pten^pc-/-^* mice, or prostate tissue harvested from 6 month old *Pkm1*;*Pten^pc-/-^* and *Pten^pc-/-^* mice as indicated. (*Pten^pc-/-^*, n=10; *Pkm2*;*Pten^pc-/-^*, n=6; and *Pkm1*;*Pten^pc-/-^*, n=6). The p value is shown for each significant difference in metabolite levels.

